# Many rare genetic variants have unrecognized large-effect disruptions to exon recognition

**DOI:** 10.1101/199927

**Authors:** Rocky Cheung, Kimberly D. Insigne, David Yao, Christina P. Burghard, Eric M. Jones, Daniel B. Goodman, Sriram Kosuri

## Abstract

Any individual’s genome contains ∼4-5 million genetic variants that differ from reference, and understanding how these variants give rise to trait diversity and disease susceptibility is a central goal of human genetics^1^. A vast majority (96-99%) of an individual’s variants are common, though at a population level the overwhelming majority of variants are rare^2–5^. Because of their scarcity in an individual’s genome, rare variants that play important roles in complex traits are likely to have large functional effects^6,7^. Mutations that cause an exon to be skipped can have severe functional consequences on gene function, and many known disease-causing mutations reduce or eliminate exon recognition^8^. Here we explore the extent to which rare genetic variation in humans results in near complete loss of exon recognition. We developed a Multiplexed Functional Assay of Splicing using Sort-seq (MFASS) that allows us to measure exon inclusion in thousands of human exons and surrounding intronic sequence simultaneously. We assayed 27,733 extant variants in the Exome Aggregation Consortium (ExAC)^9^ within or adjacent to 2,339 human exons, and found that 3.8% (1,050) of the variants, almost all of which were extremely rare, led to large-effect defects in exon recognition. Importantly, we find that 83% of these splice-disrupting variants (SDVs) are located outside of canonical splice sites, are distributed evenly across distinct exonic and intronic regions, and are difficult to predict *a priori*. Our results indicate that loss of exon recognition is an important and underappreciated means by which rare variants exert large functional effects, and that MFASS enables their empirical assessment for splicing defects at scale.

## Main Text

Common variants in a population usually contribute small, additive effects towards complex traits, as negative selection has removed large-effect deleterious alleles^10^. Human population expansion ∼10,000 years ago has left humans with an abundance of rare variation, and most Mendelian disease traits are caused by rare alleles with large effect sizes^11^. For complex traits, traditional population or computational genomic methods cannot reliably estimate the contribution of extremely rare variants, many of which have the largest effect sizes^12^. Recent whole genome and transcriptome sequencing studies of large cohorts indicate that rare variation is playing an important role in shaping global gene expression^13–15^. If rare variants are playing a large role in global gene expression and complex traits more generally, then they likely have large effect sizes due to their relative scarcity in an individual’s genome. However, new comprehensive reverse-genetic studies indicate that individual mutations in promoter and enhancer regions rarely have large effects^16–20^, which may be the result of functional redundancy between transcriptional control elements^21–23^. How can individual rare variants be broadly shaping gene expression, but at the same time rarely having large effects on transcriptional control? We can expect the mutational profiles of large-effect rare variants to mirror those that cause Mendelian traits, which are dominated by non-synonymous exonic mutations, structural and copy number variants, or mutations that affect splicing^24,25^. While copy number changes and non-synonymous mutations are easy to detect, splicing changes are more difficult to diagnose, as only mutations at canonical splice sites are easy to predict and interpret^26^.

There are several lines of evidence now accumulating that genetic variation influences traits through their effects on splicing more than previously appreciated. For common variants, large-cohort RNA-Seq studies that examine splicing are finding many splicing quantitative trait loci (sQTL), especially when considering exon-level expression differences^27–29^. Moreover, a majority of eQTLs tend to act on an individual exon level rather than the gene level, indicating that cis-eQTLs might be broadly affecting exon recognition^30^. In addition, functional genomic measurements of GEUVADIS individuals indicate that common genetic variation influencing splicing is a primary mechanism that confers susceptibility to common diseases^31^. For rare variation, analysis of bottlenecked populations find that many rare variants which segregate with large-effect expression changes are enriched at splice sites^32^. In addition, prospective transcriptional profiling studies for Mendelian diseases are increasingly finding many rare variants that affect splicing that were difficult to predict *a priori*^*33,34*^. More broadly, computational splicing predictors trained on RNA-Seq data and sequence features seem to indicate that many rare and disease variants are predicted to influence splicing levels^35^. Finally, a large-scale functional assay examining ∼5000 exonic disease mutations indicate that ∼10% of them have some effect on splicing^36^, but many functional mutations are not located close to the splice sites, suggesting that many splicing defects are likely yet to be discovered.

We developed MFASS to better understand the extent to which mutations within exons and introns can lead to large-effect exon-recognition defects. In humans, due to long intron lengths, exons are first recognized by the splicing machinery in a process called exon definition, and thus mutations that affect exon recognition often result in exon skipping rather than intron retention^8^. MFASS uses a set of three-exon, two-intron reporters in which skipping of the middle exon leads to reconstitution of fluorescence (**Fig. 1A, Supplementary Fig. 1, Supplementary Note 1**). We cloned libraries of microarray-derived oligonucleotides^37^ that encoded human exons and surrounding intronic sequences into these reporters *en masse* to make reporter libraries. These libraries are then integrated into HEK293T human cell lines using serine integrase-based site-specific integration into the AAVS1 locus (**Supplementary Fig. 6**), ensuring one copy of library sequence per cell and mitigating dosage-related issues in splicing behavior during transient tests (**Supplementary Note 2**). We used fluorescence-activated cell sorting (FACS) to separate the pooled sequence library of splicing reporters into three to four bins, which corresponds to splicing behavior ranging from exon skipping to exon inclusion. We expanded these sorted bins over several passages and observed that the sorted populations remained stable (**Fig. 1B**). We also performed bulk RT-PCR for each bin, and found that the observed RNA splicing efficiencies corresponded almost directly with observed fluorescence of the bins (**Fig. 1C**). Individual controls sorted from the library showed consistent behavior between inclusion rates estimated by RT-PCR and fluorescence output (**Supplementary Note 2**). We calculated an exon inclusion index for each sequence based on a weighted average of normalized read counts for each bin multiplied by the average exon inclusion level for that bin (**Supplementary Fig. 8A, Supplementary Methods**).

**Figure 1:**
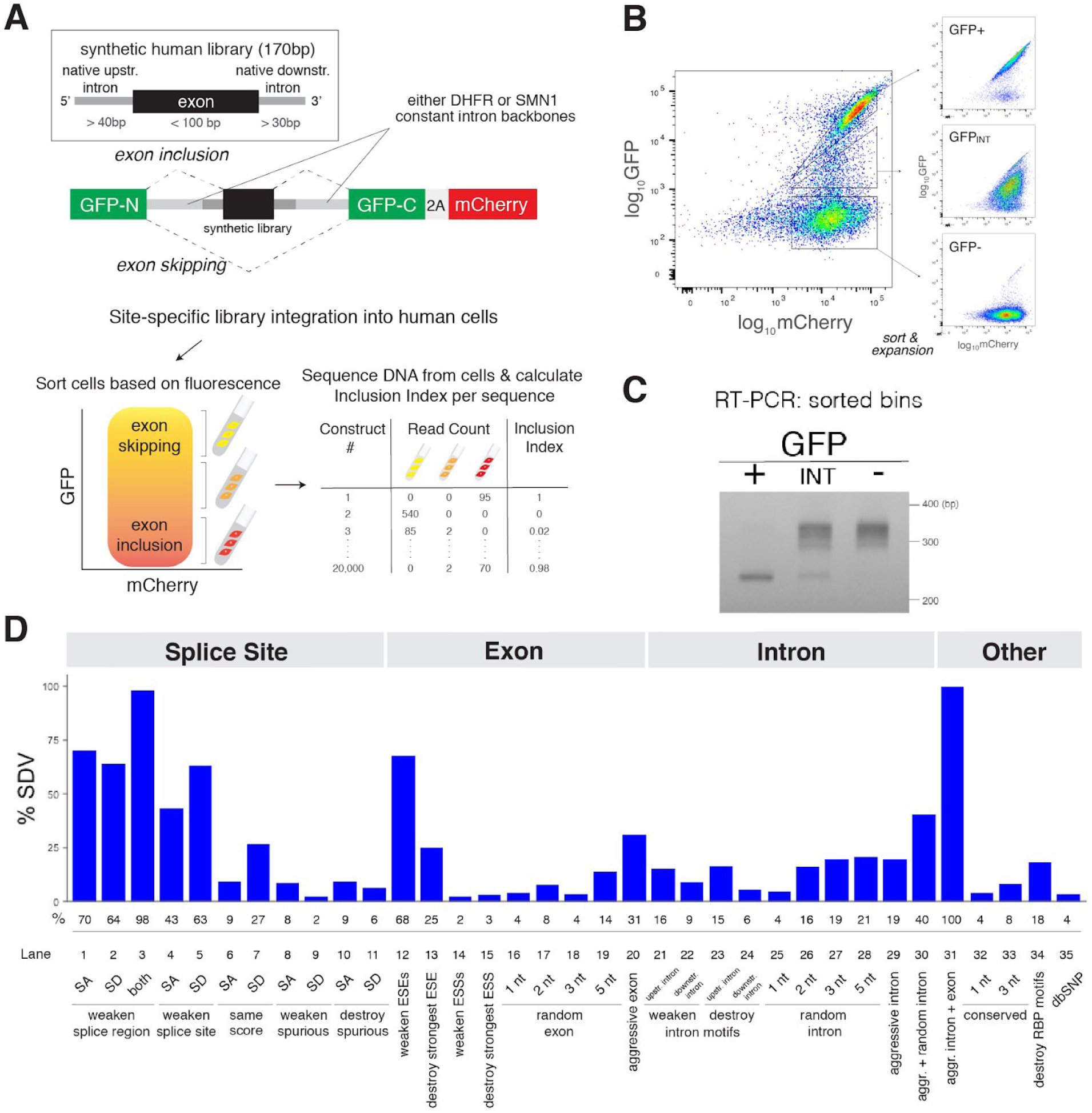
Multiplexed Functional Assay of Splicing by Sort-seq (MFASS). A. We cloned synthetic human exons (*black*) and surrounding intronic sequences (*dark grey*) into our reporter plasmid containing a split-GFP reporter with flanking constant intron backbones (*light grey*), followed by site-specific integration into HEK293T cells using Bxb1 integrase. Cells are sorted into bins based on fluorescence, followed by amplicon sequencing from cells in each sorted bin. We calculated exon inclusion index for each sequence based on a weighted average of normalized read counts across all bins multiplied by the average exon inclusion levels measured by fluorescence. **B.** We used FACS to sort the genomically-integrated SRE library into three separate populations (*left*). After expansion, the sorted populations remained stable (*right*). **C.** The observed RNA splicing efficiencies of the sorted bins as measured by RT-PCR correspond directly with observed fluorescence of the bins. **D.** Quantitative measures of exon inclusion for iteratively-designed mutations across 35 categories of splicing regulatory elements (*n* = 6713), as defined in **Supplementary Note 3**. Splice-disrupting variants (SDVs) are defined as a mutation to a wild-type exon with an inclusion index of ≥ 0.50, that is reduced by an absolute value of at least 0.5, and percentage of SDVs is indicated below each class. ESE, exonic splicing enhancer. ESS, exonic splicing suppressor. RBP, RNA-binding protein. SA, splice acceptor. SD, splice donor.

To test and validate MFASS, we first designed, built and assayed a test library of 6714 mutations aimed at perturbing regulatory elements across a randomly chosen library of 205 natural in-frame human exons and surrounding intronic sequences (Splicing Regulatory Element library, see **Supplementary Note 3**). Most library sequences are represented predominantly in one bin, showing either complete exon inclusion or skipping (**Supplementary Fig. 8A**). We tested these libraries across two constant intron backbones (SMN1 and DHFR), and found that exon inclusion metrics are highly reproducible within the backbone across biological replicates (**Supplementary Fig. 8B and C**) (*r*_*t*_ = 1.00, *P* < 10^−16^, tetrachoric; *r* = 0.94, *P* < 10^−16^, Pearson, DHFR intron backbone; *r*_*t*_ = 0.97, *P* < 10^−16^, tetrachoric, *r* = 0.89, *P* < 10^−16^, Pearson, SMN1 intron backbone), and between backbones (**Supplementary Fig. 8D**) (*r*_*t*_ = 0.96, *P* < 10^−16^, tetrachoric; *r* = 0.85, *P* < 10^−16^, Pearson). To focus on the mechanisms by which large-effect splicing changes can occur, we quantify the percentage of splice-disrupting variants (SDVs), which we define as a mutation to a wild-type exon with an inclusion index of ≥ 0.5, that is reduced by an absolute value of at least 0.5 (**Supplementary Note 4**).

We find that mutations within the splice sites tend to cause the most SDVs (**Fig. 1D**). Mutations intended to weaken these sites individually result in SDVs 50-75% of the time (lanes 1,2,4,5), and 98% of the time when mutating both donor and acceptor simultaneously (lane 3). This is likely an underestimate as mutations eliminating splice site recognition may be utilizing alternative splice acceptors or donors, which cannot be distinguished from exon inclusion by MFASS. Within exons, mutations can still have strong effects. Encoded synonymous mutations intended to weaken previously identified exonic splicing enhancers lead to SDVs ∼60% of the time (lane 12), and removing the strongest identified ESE alone results in 25% SDVs (lane 13). More generally, existing splicing metrics such as MaxEnt for splice site strength (**Supplementary Fig. 11A and B**) or exon hexamer metrics (**Supplementary Fig. 11C and D**) are consistent with predicted effects on splicing behavior as evidenced by the change in inclusion index, albeit these metrics often do not provide much predictive value.

While these results indicate that mutations intended to alter previously recognized motifs can commonly lead to loss of exon recognition, we wanted to explore the extent to which natural genetic variation result in SDVs. We designed and synthesized all catalogued exonic and intronic single nucleotide variants (SNVs) from the Exome Aggregation Consortium (ExAC) for 2902 human exons, and quantified the effects of more than half of these SNVs found across 2339 exons (52.4%, 27,733/52,965) (**Supplementary Note 5**). Overall, we found that 3.8% (1050/27,733) of ExAC variants led to almost complete loss of exon recognition (**Fig. 2A**). We observe almost equal contributions of SDVs from introns (53%) and exons (47%) (**Fig. 2A**), broadly spread across 543 human exon backgrounds (**Supplementary Fig. 13A**). We found that 68% of splice site variants are SDVs (**Fig. 2B, left**), again noting that alternative 5’ and 3’ splice site usage are measured as false negatives for MFASS. Compared with splice site variants, variants in the broader splice region, synonymous exonic variants, non-synonymous exonic variants, and deeper intronic variants disrupt splicing more rarely at 8.5%, 3.0%, 3.1%, and 1.5% respectively (**Fig. 2B, left**). However, because SNVs are not equally distributed amongst these categories, splice site SDVs only constitute 17% of all SDVs, whereas intron variants, which are the least sensitive to splicing disruption, comprised 19% of SDVs (**Fig. 2B, right**). SNVs at the splice sites are rare in our library (**Fig. 2C, bottom panel line 3**), and also for all ∼7.4 million ExAC variants (**Supplementary Fig. 14**). The larger number of variants in regions away from the splice sites outweighs their reduced sensitivity (**Fig. 2C**), and contribute 83% of the 1050 SDVs reported here.

**Figure 2:**
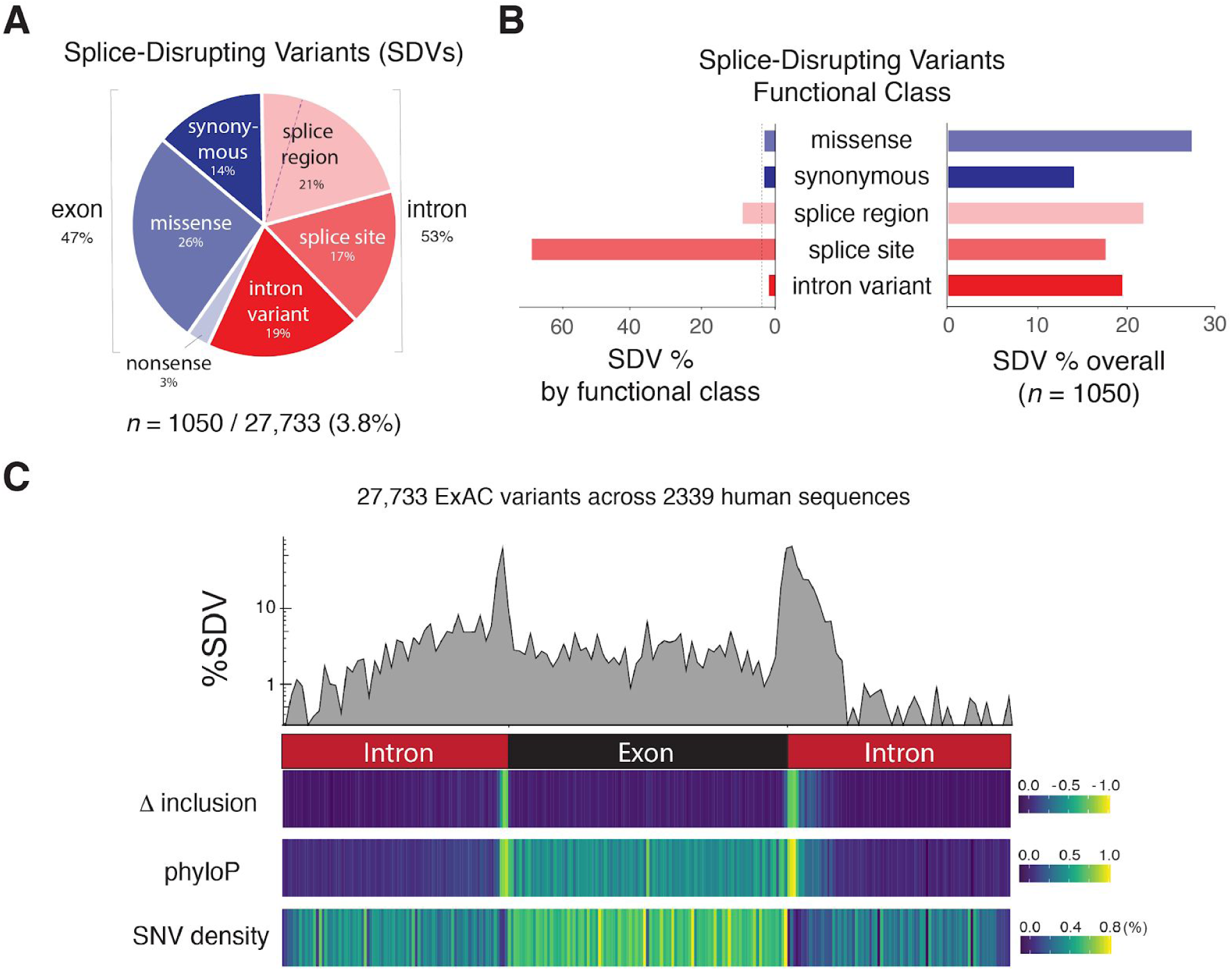
Characterization of SDVs amongst 27,733 ExAC SNVs in or near 2339 human exons. A. SDVs (*n =* 1050) are split almost equally between exonic and intronic regions (*blue* and *red* respectively). Splice region variants that fall within exonic regions (4%) and intronic regions (17%) are separated by a dashed line. **B.** Splice site mutations are by far the most likely region to result in an SDV (*left*). However, because SNVs at splice sites are relatively rare, SDVs in regions other than the splice site constitute 83% of all SDVs (*right*). **C.** The percentage of SDVs as a function of position along the exon and surrounding intron sequence shows that splice donor regions are more sensitive than splice acceptor regions (*top panel*). We also plot the change the average change exon inclusion index (δinclusion index), mammalian evolutionary conservation (phyloP score averages), and the ExAC SNV density as a function of location. Each bin corresponds to 1-2 nucleotide per position, and locations are relative as we test a range of exon lengths.

A number of population genetic, evolutionary, and functional characterizations indicate that our measured SDVs are relevant. *First*, the proportion of SNVs that are SDVs shows significant reductions as a function of allele frequency (chi-squared test, *P* = 1.12 x 10^−3^). Consistent with population genetic theory, a vast majority of our SDVs are extremely rare (**Fig. 3A**). *Second*, we find a significantly lower SDV rate (∼2x) within genes that rarely have protein-truncating mutations within ExAC, indicating strong functional constraint (pLI ≥ 0.9)^9^ (**Fig. 3B**) (two-tailed Fisher’s exact test, *P* = 1.30 x 10^−12^). *Third*, SNVs that are SDVs show significantly stronger evolutionary constraint, suggesting purifying selection at these sites (Student’s *t* test, *P* < 10^−16^) (**Fig. 3C**). *Fourth*, nucleotide positions under strong evolutionary constraint have higher rates of SDVs, and this is especially apparent within introns (two-tailed Fisher’s exact test, *P* < 10^−16^) (**Fig. 3D**). However, this conservation has limited predictive power, because within introns there are many more SNVs at neutral sites than sites under strong constraint, and within exons most sites are highly conserved (**Fig. 3E**). *Fifth*, for exonic SNVs, we observed that SDVs significantly reduce exon hexamer scores when compared with non-SDVs, suggesting that SDVs are disrupting important functional sites for exon recognition (Student’s *t* test, *P* < 10^−16^) (**Fig. 3F**). *Sixth*, motif enrichments at the splice acceptor suggests that SDVs enriched for T to A mutations disrupt the area near the mechanistically important polypyrimidine tract, while for splice donors we find that guanine-rich motifs are less tolerated (**Supplementary Fig. 13C**). *Sixth*, we verified SDVs individually in transient expression assays across multiple functional categories (**Fig. 2B**), and found that 9/11 showed large-effect splicing defects, with all 11 showing reduced exon inclusion as compared to their respective wild-type sequences (**Supplementary Fig. 15**). We also tested the effect of longer intronic contexts on detected SDVs, and found that 17/23 SDVs showed large defects in splicing, with only 1/23 mutations showing no appreciable exon recognition defect (**Supplementary Fig. 15**).

**Figure 3:**
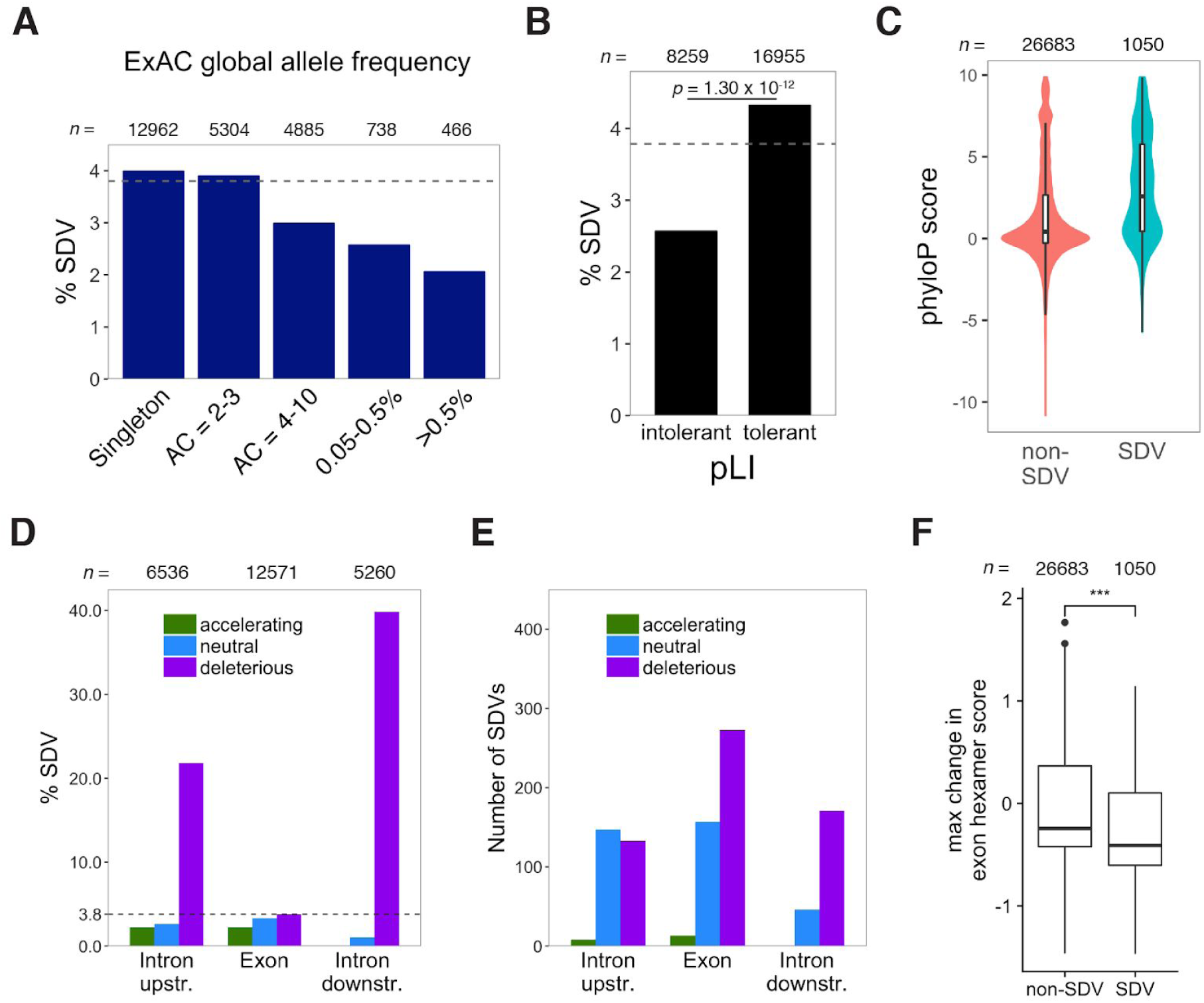
Characteristics of Splice-Disrupting Variants (SDVs). A. The percentage of SDVs as a function of allele frequency (AC, allele count) shows significant reductions across allele frequencies (chi-squared test, *P* = 1.12 x 10^−3^). Almost all (98.1%) of the ExAC variants assayed were rare (global MAF <0.5%). **B.** We observe significantly fewer SDVs for exons within those genes that are predicted to be intolerant to loss-of-function alleles (pLI ≥ 0.9) (two-tailed Fisher’s exact test, *P* = 1.30 x 10^−12^), with the overall percentage shown as a dashed line (3.8%). **C.** SDVs are under stronger evolutionary constraint as evidenced by higher overall phyloP scores (Student’s *t* test, *P* < 10^−16^). **D.** Within introns, we find that positions that are evolutionarily constrained (deleterious, phyloP > 2.0) have a higher SDV rate than those under neutral (−1.2 ≤ phyloP ≤ 1.2) or accelerating selection (phyloP < −2.0) (two-tailed Fisher’s exact test, *P* < 10^−16^). **E.** Because there are more SNVs outside of regions of high intron conservation, there are still many SDVs located within sites displaying neutral selection. **F.** We observed a significantly lower maximum change in predicted exonic hexamer score within exonic SDVs than non-SDVs (Student’s *t* test, *P* < 10^−16^).

Our results indicate that traditional metrics for assessing how mutations affect splicing are likely to fail, because while it is known that splice site variants are likely deleterious, it has been unclear to what extent rare genetic variation affects splicing outside of these sites. For example, existing variant effect predictors for missense mutations, such as Polyphen and SIFT, either largely provide no annotation for SDVs or call them benign (**Fig. 4A**). Moreover, the SDV rate in synonymous mutations, which are usually assumed to be benign, is nearly equivalent to missense variants. We used a number of contemporary variant effect predictors that are capable of predicting the effects of non-coding variation based on both functional genomic and/or evolutionary information (CADD^38^, DANN^39^, FATHMM-MKL^40^, fitCons^41^, LINSIGHT^42^, phastCons^43^ and phyloP^44^), as well as two specifically designed for splicing (SPANR^35^ and HAL^45^) (**Fig. 4B**). Most predictors have low precision, with several providing no better prediction than random guessing. FATHMM-MKL, CADD, and DANN perform best amongst those not trained specifically for splicing, but only achieve ∼7-8% precision at any appreciable recall. Much of their power is the result of the ability to call intronic SDVs (**Supplementary Fig. 16A and B**), likely due to their increased conservation. Not surprisingly, those predictors trained specifically for calling splice defects perform best. At equivalent effect size compared to our assay (>50% splicing disruption), SPANR achieves 44.5% precision, though only a minority of the SDVs are called (11.8%) (**Fig. 4B**). As we lower the threshold for calling an SDV (i.e., the predicted effect size of an SNV), SPANR can achieve 14.9% precision at 50% recall level, though the predicted effect size is ∼2% loss of inclusion. More generally, SPANR effect sizes poorly predict our observed inclusion rates (*R*^*2*^ = 0.11, **Supplementary Fig. 16C**). The increased power of SPANR over other predictions is largely due to its ability to predict exonic SDVs. HAL provides even better precision in these exonic regions (**Supplementary Fig. 16B**), but only calls SNVs within exons.

**Figure 4:**
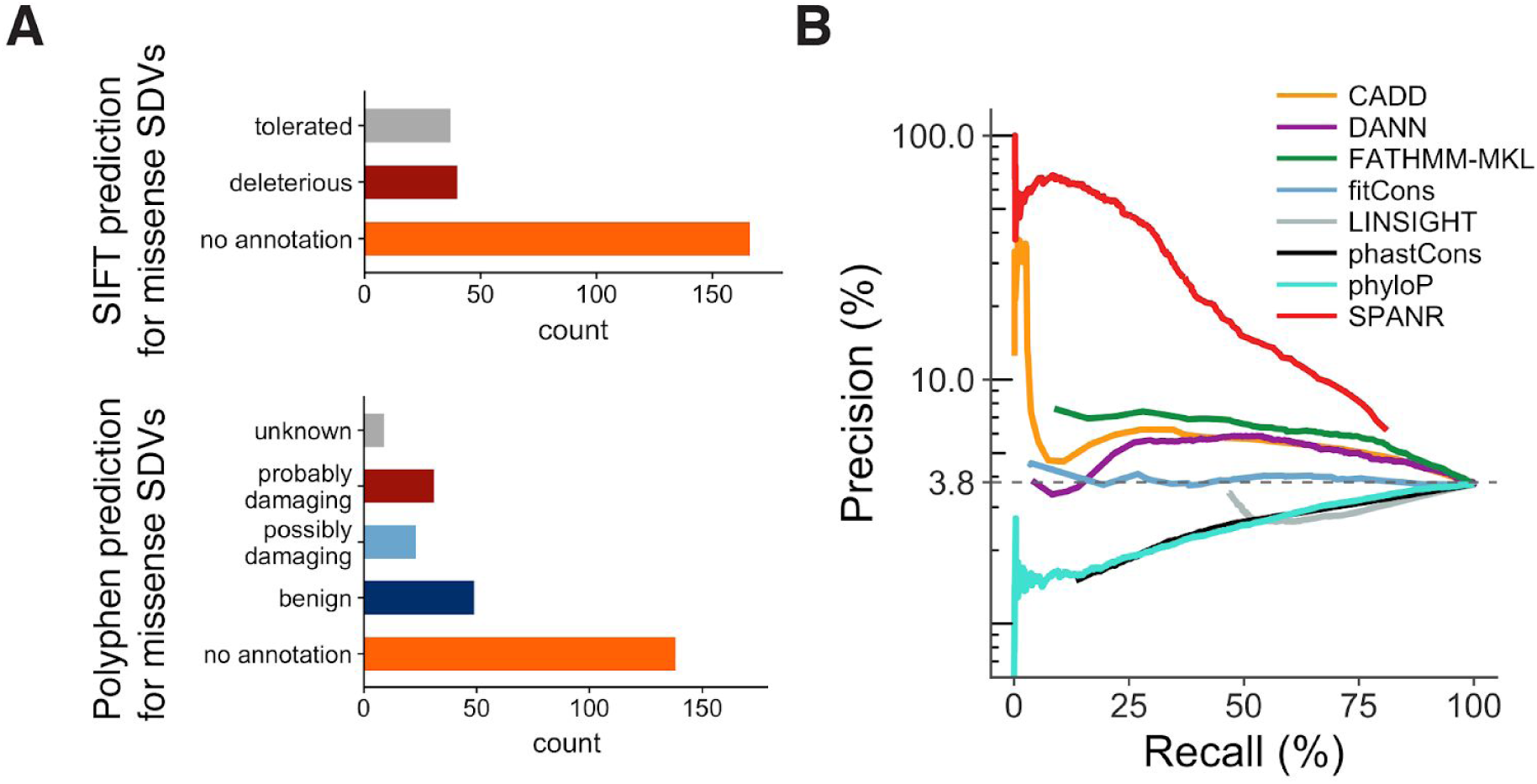
Evaluation of genomic and deep-learning predictors for rare genetic variants on splicing. A. Functional prediction from SIFT and Polyphen for missense SDVs show few are predicted to be loss of function. **B.** Precision-recall curves for algorithms that can predict splicing or non-coding genetic variants. Dashed line in **B** indicates the overall percentage of SDVs (3.8%, 1050/27,733) from the MFASS assay.

In this work, we test over half of the variants found in 2339 human exons across ∼60,000 individuals and observed that 3.8% of these variants (1050/27,733) can cause loss of exon recognition. There are a number of technical and biological reasons we may be over- or under-estimating the number of SDVs by using MFASS, including selection bias in the exons we chose, ascertainment bias in our experimental workflow, as well as limitations in the reporter assay and the choice of cell line (**Supplementary Note 6**). The rate of SDVs we find here is surprisingly high. It is ∼73% of the rate of probably damaging variants predicted by PolyPhen for the same set of SNVs (5.2%, 1437/27,733), and ∼3-fold higher than the observed rate of protein truncating variants found in ExAC as a whole (1.3%, 121,309/7,404,909)^9^. Importantly, we show that most of the SDVs would not be easily recognized, as only 17% of the SDVs we found are in canonical splice sites. We would expect such exon skipping events to be highly detrimental to not only protein function, but if our results generalize to exons that do not preserve frame, can cause large changes to mRNA stability through nonsense-mediated mRNA decay^46^. This may help explain why extremely rare variation seems to have large predicted effects on gene expression even though we rarely see individual mutations having large effects on transcriptional control elements^15^. Compared to other multiplexed splicing reporters^36,45,47–49^, MFASS is unique in that it screens both exonic and intronic variants, provides increased power for detecting large-effect loss-of-function variants, uses long constant intron backbones, and site-specifically integrates reporters into the same safe-harbor loci at single copy. MFASS is best suited for screening large numbers of large-effect rare variants, and when used as such can likely be scaled by several orders of magnitude. More broadly, MFASS combined with other assays of variant effects^50^ can help interpret variants found in large exome datasets to get a broader understanding for how rare, *de novo*, and somatic variants are shaping complex traits and diseases^51^.

## Acknowledgments

This work was supported by the National Institutes of Health (5U01HG007912 & DP2GM114829 to S.K.), the NIH Biomedical Big Data Training Grant (T32CA201160 to C.B.), Searle Scholars Program [to S.K.], Department of Energy (DE-FC02-02ER63421 to S.K.), UCLA, and Linda and Fred Wudl. We thank Ron Weiss for the original landing pad cell line, Felicia Codrea and Jessica Scholes (UCLA BSCRC flow cytometry core), and the BSCRC high throughput sequencing core for technical assistance. We thank X. Grace Xiao, Douglas Black, and George Church for guidance while developing MFASS.

## METHODS

### Microarray-derived oligonucleotide library design

We obtained microarray-derived oligonucleotides of 200 to 212 bp from Agilent Technologies to generate synthetic DNA libraries. We selected human exons that are less than 100 bp and begin and end on frame 0 from the Ensembl mySQL server^54^ (Ensembl release 73, hg19 assembly). We designed a 170-bp intron-exon-intron sequence library *in silico* containing all human exons fulfilling above criteria, which includes at least 40 bp of upstream intron and at least 30 bp of downstream intron (*n* = 9634), with the exon in the middle. We added extra native intronic sequences as length limitations allowed (i.e., if exons were shorter), split between the upstream and downstream equally with an extra base added to the donor side for odd number of bases added.

For the SRE library, we randomly chose 230 exons from our wild-type library, and computationally designed 60-80 synonymous mutations per sequence using a custom software we developed (**Supplementary Note 3**), that correspond to specific functional classes of regulatory elements governing splicing. Finally, a pair of 15-mer amplification primer sequences, containing AscI/PacI restriction sites, were added to yield 200-mer sequences for DNA synthesis. For the SNV library, we used a library of 2902 exons that showed high inclusion using MFASS (**Supplementary Note 5**). We obtained single nucleotide variants (SNVs) from the Exome Aggregation Consortium^55^ (ExAC, version 0.3.1). We stored hg19 genomic coordinates of each sequence in BED file format, and used bcftools to intersect the ExAC variants with our library of wild-type human exons to subset all relevant SNVs. We only synthesized variants with a filter status of “PASS”, and generated all alternate alleles (up to 3) if more than one alternate allele was indicated. These sequences were filtered to (i) exclude sequences containing unique NheI or AgeI restriction sites used for library cloning and (ii) include SNVs only within nucleotides 11 through 160 of each 170 bp library sequence to avoid possible spurious interactions with restriction sites. Finally, a pair of 15-mer amplification primer sequences, as well as NheI/AgeI restriction sites, were added to yield 212-mer sequences for DNA synthesis (**Supplementary Note 5**).

### Library amplification and cloning

Oligonucleotide libraries were amplified with KAPA HiFi HotStart (Kapa Biosystems) using 500 pg of oligonucleotide library using biotinylated primers flanking the human exon libraries. For the SRE library, each sublibrary was purified and digested with AscI and PacI (New England Biolabs) at 37°C to cleave off the priming sites. The resulting ends were removed by M-270 streptavidin beads (Invitrogen) and the supernatant was collected. For the SNV library, we performed similar procedures as above with the following alterations: we performed emulsion PCR, and processed the amplicons with NheI and AgeI (New England Biolabs) at 37°C before ligation-based cloning and transformation into electrocompetent *E. coli* (New England Biolabs) (**Supplementary Methods**).

### Generation of landing pad cell line

For site-specific integration of exon libraries in HEK293T cells, we engineered a chromosomal landing pad cell line, which allows stable expression of splicing reporter library at the AAVS1 locus, which is modified from Duportet et al.^56^ by CRISPR-Cas9 in order to remove expression of the endogenous YFP gene. We characterized 25 clones expanded from single cells by flow cytometry, microscopy and genomic PCR, and selected a clone (which we termed RCA7) that does not express any YFP or RFP fluorescence for our current study.

### Serine-integrase based genomic integration of synthetic libraries

We prepared reporter plasmids for mammalian cell transfection and generated site-specific, genome-integrated reporter cell libraries. For splicing reporter library experiments, after genome integration and puromycin selection, each biological replicate represented ∼200-fold library coverage. For all cell libraries, landing pad cells were transfected with the library and Bxb1 serine integrase for 72 hours (4:1 ratio), and then selected with 5 μg/mL puromycin (Life Technologies). Cells were selected for integrants and subsequently passaged serially for at least 18 days before cell sorting (**Supplementary Methods**).

### Fluorescence-activated cell sorting

Cells harboring variant libraries were sorted using a FACSAria III (BD Biosciences) into bins based on GFP fluorescence, given a minimal amount of mCherry fluorescence. We eliminated dead cells, debris, and doublets based on forward and side scatter, and single-color and double-negative controls were used for gating and calibration. We sorted based on GFP and mCherry fluorescence for the SRE library and SNV library version 1 (3 bins) or SNV library version 2 (4 bins), for roughly 2-10 million cells per bin that is proportional to bin size. Sorted sub-libraries for each replicate were grown separately and passaged. For the SRE libraries, we sorted cells into 3 bins (**Fig. 1B**). For the SNV library version 1, we performed two sequential cell sorts to obtain the reporter libraries (**Supplementary Fig. 4**). For the SNV library version 2, we sorted cells based on GFP fluorescence into four bins (**Supplementary Fig. 5**). We obtained sorts for two biological replicates for all these libraries.

### DNA-Seq of FACS-sorted libraries

We extracted genomic DNA from 10-20 million cells for the three to four sorted populations using Qiagen blood and cell culture DNA midi kit (Qiagen). For the SRE library, we amplified each sublibrary for ∼300-fold amplicon coverage, and reactions were performed in 96-well format in three to nine 50 μL reactions for each sorted bin proportional to the number of cells sorted. Per biological replicate, we amplified library variants from genomic DNA with KAPA HiFi HotStart (Kapa Biosystems), using 2-5 μg of template. PCR primers were designed to give 300-600 bp amplicons, which we subsequently attach Illumina adapters by a secondary amplification necessary for next-generation sequencing (**Supplementary Methods**). The amplicons were gel-extracted on 1% agarose gel and quantified using Agilent Tapestation 2200. For the SNV library, sorted libraries were indexed by PCR amplification, in twenty-four 50 μL reactions for GFP_neg_ and eight 50 μL for all other sublibraries, with the same 2-5μg per reaction of genomic DNA.

### Reverse transcription-PCR

RNA from sorted sub-libraries as well as individual control exons were extracted using Qiagen RNEasy MiniKit. Reverse transcriptions were performed using Superscript IV (Thermo Fisher Scientific) according to manufacturer’s protocol, which primes to a region in emerald GFP exon 2 (**Supplementary Methods**).

### DNA-Seq read processing and filtering

SRE library MFASS DNA-Seq datasets were generated from two Illumina MiSeq 300-bp paired-end sequencing runs and one Illumina HiSeq 2500 150-bp paired-end sequencing run. SNV library (version 1) MFASS DNA-Seq dataset was generated from Illumina MiSeq 300-bp paired-end sequencing. SNV library (version 2) MFASS DNA-Seq dataset was generated from Illumina NextSeq 2500 150-bp paired-end sequencing. We removed read pairs with any ambiguous “N” base calls, followed by read pair merging with *bbmerge* from the BBMap suite^57^ (BBtools package version 37). We developed custom Python and bash scripts to filter for perfect reads aligned to our reference sequences, from which we can aggregate read counts for sequences from each sorted bin. We then further process these read counts to calculate inclusion index (see below section on the quantification of inclusion index).

We applied minimal sequencing depth filters of at least 5 reads across all bins for the SRE library. Our SRE library size was 16,717 (5975 wild-type sequences, 10,683 mutants, 59 controls) for the SMN1 intron backbone, and 13,922 (4920 wild-type sequences, 8942 mutants, 60 controls) for the DHFR intron backbone. For functional analyses, we required that the index agrees within 0.30 across the DHFR and SMN1 intron backbones, resulting in a library size of 10,482 (3714 wild-type sequences, 6713 mutants, 55 controls). For the SNV library, we only analyzed a mutant sequence if its corresponding wild-type sequence has an inclusion index of ≥ 0.5. We observed 43,398 mutants (version 2) appearing at least once in either replicate that fulfil above criteria. For downstream analyses of the SNV library, we applied sequencing depth filters of at least 10 reads across all bins and inclusion indice filters for biological replicates within 0.20. Our SNV library size (version 1) is 6768 (1981 wild-type sequences, 3853 mutants, 934 controls). Our SNV library size (version 2) is 31,144 (2339 wild-type sequences, 27,733 mutants, 1072 controls).

### Population genetic data analysis

Annotation of variants for individual human samples in VCF format were obtained from the Exome Aggregation Consortium^9^ (ExAC, version 0.3.1), including global allele frequencies. We binned ExAC global allele frequency similar to the ExAC study, and tested for significant difference between allele frequency bins using chi-squared test of independence. We also obtained gene level evolutionary constraint estimates from ExAC based on probability of loss-of-function intolerance (pLI), and defined genes that are extremely intolerant of loss-of-function as those with a pLI score ≥ 0.9. We then tested for genes with enrichment in splice-disrupting variants (SDVs) using Fisher’s exact test.

### Functional genomic analysis of SNVs

We classified our variants using the Ensembl Variant Effect Predictor^58^ (VEP v80), and filtered the most severe sequence ontology term for a given variant. We obtained phyloP 100-way (v1.4) nucleotide conservation for the hg19 genome for the SNV library, and classified quickly evolving regions of the genome (accelerating, phyloP < −2.0), neutral selection (−1.2 ≤ phyloP ≤ 1.2) and highly conserved region of the genome (deleterious, phyloP > 2.0).

To compute genome-wide locations of ExAC SNVs by gene regions, we used GENCODE^59^ (release 27, hg38 assembly) for exon annotation, and bedtools^60^ to annotate intronic regions by subtracting exon coordinates from gene coordinates. To determine the density of SNVs for each genomic position, we determined the number of SNVs averaged at each relative scaled position for the SNV library as well as genome-wide SNVs from the ExAC consortium. In particular, we calculate scaled positions for each SNV to normalize for variable intron and exon lengths. Relative position is set such that the boundary of upstream intron/5’ exon = 0, and the boundary of 3’ exon/downstream intron boundary = 1. Splice site variants are defined as 2 bp of intron adjacent to exon by the Variant Effect Predictor classification, whereas splice region variants are located 2 bp into the exon and 8 bp into the intron, excluding splice sites.

### Quantification of exon inclusion from Sort-seq

We normalized bin counts based on read depth (reads per million, RPM) and corresponding bin population percentage after FACS using the following formula:

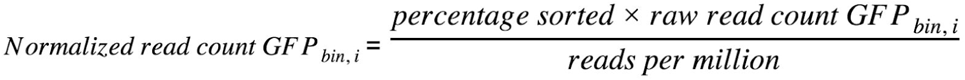

We calculated exon inclusion index for each sequence based on a weighted average of normalized counts across all bins. Bin weights are assigned proportionally based on GFP fluorescence measurements of individual bins that correspond to the extent of exon inclusion or skipping. For the splicing regulatory element (SRE) library and single nucleotide variant (SNV) library, version 1:

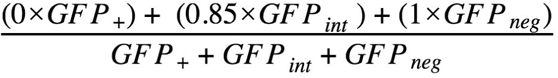

For the SNV library, version 2:

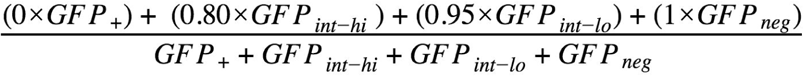

The change in inclusion index for an individual library sequence between wild-type (WT) and mutant is computed as follows:

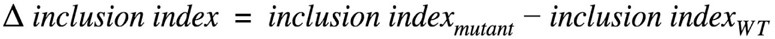

A positive δinclusion index denotes increased exon inclusion for the mutant relative to WT, while a negative δinclusion index denotes increased exon skipping for the mutant relative to WT.

### *k* -mer LOGO motif enrichment analysis

To define potential disruption of *k*-mer motifs by ExAC SNVs, we performed *k*-mer based motif enrichment analysis using *k*pLogo^61^ for both 3’ splice site/region (−20 to +3 positions, left intron/exon junction in all schematics) and 5’ splice site/region (−3 to +6 positions, right intron/exon junction in all schematics), using a p-value cutoff of *P* < 0.01, gapped *k*-mer length of *k* = 1,2,3,4 and fixation frequency of 0.75. Based on our MFASS dataset, SDVs are background-corrected against non-SDVs to obtain motif logos that are enriched or depleted at each nucleotide.

### Exon hexamer score analysis

We implemented the hexamer additive linear (HAL^45^) model, which estimates a splicing strength score for every possible exon hexamer. A positive score indicates the hexamer is more likely to activate nearby splice sites, and a negative score indicates the hexamer is more likely to silence nearby splice sites. For each variant, we calculated the change in score at each position relative to the wild-type sequence, and identified the maximum change in score. We compared the distribution of maximum score change between SDVs and non-SDVs using the two-sample Student’s *t-*test.

### Implementation of HAL model for exonic SNVs

To evaluate predictive performance of exonic hexamers on exonic SNVs, we adapted the HAL model to compute δψ for our sequences based on exon hexamer scores. The large training set enabled a general model of hexamer strength in exons but not introns, and therefore its performance is only assessed for exon variants.

### Assessment of external variant prediction algorithms

To computationally predict the effects of rare genetic variants on splicing, we used various prediction algorithms that are able to assess coding and/or non-coding SNVs in our assay. For the purpose of method comparison, we selected δinclusion index ≤ −0.50 as the threshold for splice-disrupting variant (SDV) and designate our calls as true positives. To evaluate a more general predictive SNV model on both exonic and intronic single nucleotide variants, we obtained features for the changes in percent-spliced-in (δψ) for the SNV library across the genome from SPANR^62^ (splicing-based analysis of variants). Performance is assessed by varying the δinclusion index threshold at which a variant is called splice-disrupting.

We obtained features for five genomic predictors based on the hg19 assembly: raw CADD scores from CADD version 1.3 (r0.3 Exome Aggregation Consortium dataset), DANN whole-genome SNV scores (Nov. 2014 version), FATHMM-MKL (Jan. 2015 version), fitCons multi-cell (i6 dataset) highly significant scores (*p* < ∼0.003), and LINSIGHT (Apr. 2017 version). We assessed performance by varying the score threshold at which a variant is called splice-disrupting (considering whether the score is positively or negatively correlated to δinclusion index). To consider the predictive power of conservation alone, we obtained phyloP 100-way (v1.4) nucleotide conservation for the hg19 genome for the SNV library. In addition, we obtained phastCons^63^ (v1.4) scores for 100-way eutherian mammalian nucleotide conservation for our SNV library and genome-wide SNVs from the ExAC consortium. To assess the functional effects of missense, exonic single nucleotide variants from the SNV library, we used variant annotations from PolyPhen (version 2.2.2) and SIFT (version 5.2.2).

### Assessment of deep learning and genomic predictors

We assessed above predictors using receiver operating characteristic and precision-recall analysis. We used the pROC package version 1.10.0 to compute and plot the ROC curves, calculate the 95% confidence interval, and calculate the area under the curve. The precision recall curves were plotted with a custom function which evaluates each method by varying the score threshold at which a sequence is classified as an SDV, and calculating the corresponding precision and recall. The area under the precision recall curve is calculated with the trapz function in R.

### Software

*bbmerge* from the BBMap suite^57^ (BBtools package version 37) was used to merge raw paired-end sequencing files. Custom python and bash scripts used for read processing, and mapping reference and synthetic error read counts. Further analysis was performed with Python 2.7, using Pandas v0.21.0 and Numpy v1.13.3, and R v3.4.2, using dplyr v0.7.4 and ggplot2 v2.2.1. Variant analyses were performed using Ensembl Variant Effect Predictor (v80), *k*pLogo (2017), CADD (v1.3), DANN (Nov. 2014 version), FATHMM-MKL (Jan. 2015 version), fitCons (i6 dataset), HAL (git/ca54d11), LINSIGHT (Apr. 2017 version), phastCons (v1.4), phyloP (v1.4), PolyPhen (version 2.2.2) and SIFT (version 5.2.2) and SPANR (git/5bd33c0).

### Code availability

All codes needed to reproduce the analyses is included in the following repository: https://github.com/KosuriLab/MFASS

### Data availability

Raw sequencing data are available upon publication. Processed data sets are available upon request.

## References

1. Auton, A. et al. A global reference for human genetic variation. Nature 526, 68–74 (2015).

2. Tennessen, J. A. et al. Evolution and functional impact of rare coding variation from deep sequencing of human exomes. Science 337, 64–69 (2012).

3. Nelson, M. R. et al. An abundance of rare functional variants in 202 drug target genes sequenced in 14,002 people. Science 337, 100–104 (2012).

4. UK10K Consortium et al. The UK10K project identifies rare variants in health and disease. Nature 526, 82–90 (2015).

5. Montgomery, S. B., Lappalainen, T., Gutierrez-Arcelus, M. & Dermitzakis, E. T. Rare and common regulatory variation in population-scale sequenced human genomes. PLoS Genet. 7, e1002144 (2011).

6. Bomba, L., Walter, K. & Soranzo, N. The impact of rare and low-frequency genetic variants in common disease. Genome Biol. 18, 77 (2017).

7. Gibson, G. Rare and common variants: twenty arguments. Nat. Rev. Genet. 13, 135–145 (2012).

8. Baralle, D. & Buratti, E. RNA splicing in human disease and in the clinic. Clin. Sci. 131, 355–368 (2017).

9. Lek, M. et al. Analysis of protein-coding genetic variation in 60,706 humans. Nature 536, 285–291 (2016).

10. Altshuler, D., Daly, M. J. & Lander, E. S. Genetic mapping in human disease. Science 322, 881–888 (2008).

11. Keinan, A. & Clark, A. G. Recent explosive human population growth has resulted in an excess of rare genetic variants. Science 336, 740–743 (2012).

12. Uricchio, L. H., Zaitlen, N. A., Ye, C. J., Witte, J. S. & Hernandez, R. D. Selection and explosive growth alter genetic architecture and hamper the detection of causal rare variants. Genome Res. 26, 863–873 (2016).

13. GTEx Consortium et al. Genetic effects on gene expression across human tissues. Nature 550, 204–213 (2017).

14. Li, X. et al. The impact of rare variation on gene expression across tissues. Nature 550, 239–243 (2017).

15. Hernandez, R. D. et al. Singleton Variants Dominate the Genetic Architecture of Human Gene Expression. (2017). doi:10.1101/219238

16. Canver, M. C. et al. BCL11A enhancer dissection by Cas9-mediated in situ saturating mutagenesis. Nature 527, 192–197 (2015).

17. Diao, Y. et al. A new class of temporarily phenotypic enhancers identified by CRISPR/Cas9-mediated genetic screening. Genome Res. 26, 397–405 (2016).

18. Rajagopal, N. et al. High-throughput mapping of regulatory DNA. Nat. Biotechnol. 34, 167–174 (2016).

19. Sanjana, N. E. et al. High-resolution interrogation of functional elements in the noncoding genome. Science 353, 1545–1549 (2016).

20. Gasperini, M. et al. CRISPR/Cas9-Mediated Scanning for Regulatory Elements Required for HPRT1 Expression via Thousands of Large, Programmed Genomic Deletions. Am. J. Hum. Genet. 101, 192–205 (2017).

21. Osterwalder, M. et al. Enhancer redundancy provides phenotypic robustness in mammalian development. Nature 554, 239 (2018).

22. Frankel, N. et al. Phenotypic robustness conferred by apparently redundant transcriptional enhancers. Nature 466, 490–493 (2010).

23. Hong, J.-W., Hendrix, D. A. & Levine, M. S. Shadow enhancers as a source of evolutionary novelty. Science 321, 1314 (2008).

24. Bamshad, M. J. et al. Exome sequencing as a tool for Mendelian disease gene discovery. Nat. Rev. Genet. 12, 745–755 (2011).

25. Chong, J. X. et al. The Genetic Basis of Mendelian Phenotypes: Discoveries, Challenges, and Opportunities. Am. J. Hum. Genet. 97, 199–215 (2015).

26. Jian, X., Boerwinkle, E. & Liu, X. In silico tools for splicing defect prediction: a survey from the viewpoint of end users. Genet. Med. 16, 497–503 (2014).

27. Zhang, X. et al. Identification of common genetic variants controlling transcript isoform variation in human whole blood. Nat. Genet. 47, 345–352 (2015).

28. Ongen, H. & Dermitzakis, E. T. Alternative splicing QTLs in European and African populations using Altrans, a novel method for splice junction quantification. (2015). doi:10.1101/014126

29. GTEx Consortium. Human genomics. The Genotype-Tissue Expression (GTEx) pilot analysis: multitissue gene regulation in humans. Science 348, 648–660 (2015).

30. Ramasamy, A. et al. Genetic variability in the regulation of gene expression in ten regions of the human brain. Nat. Neurosci. 17, 1418–1428 (2014).

31. Li, Y. I. et al. RNA splicing is a primary link between genetic variation and disease. Science 352, 600–604 (2016).

32. Pala, M. et al. Population-and individual-specific regulatory variation in Sardinia. Nat. Genet. 49, 700–707 (2017).

33. Cummings, B. B. et al. Improving genetic diagnosis in Mendelian disease with transcriptome sequencing. Sci. Transl. Med. 9, (2017).

34. Kremer, L. S. et al. Genetic diagnosis of Mendelian disorders via RNA sequencing. Nat. Commun. 8, 15824 (2017).

35. Xiong, H. Y. et al. RNA splicing. The human splicing code reveals new insights into the genetic determinants of disease. Science 347, 1254806 (2015).

36. Soemedi, R. et al. Pathogenic variants that alter protein code often disrupt splicing. Nat. Genet. 49, 848–855 (2017).

37. LeProust, E. M. et al. Synthesis of high-quality libraries of long (150mer) oligonucleotides by a novel depurination controlled process. Nucleic Acids Res. 38, 2522–2540 (2010).

38. Kircher, M. et al. A general framework for estimating the relative pathogenicity of human genetic variants. Nat. Genet. 46, 310–315 (2014).

39. Quang, D., Chen, Y. & Xie, X. DANN: a deep learning approach for annotating the pathogenicity of genetic variants. Bioinformatics 31, 761–763 (2015).

40. Shihab, H. A. et al. An integrative approach to predicting the functional effects of non-coding and coding sequence variation. Bioinformatics 31, 1536–1543 (2015).

41. Huang, Y.-F., Gulko, B. & Siepel, A. Fast, scalable prediction of deleterious noncoding variants from functional and population genomic data. Nat. Genet. 49, 618–624 (2017).

42. Gulko, B., Hubisz, M. J., Gronau, I. & Siepel, A. A method for calculating probabilities of fitness consequences for point mutations across the human genome. Nat. Genet. 47, 276–283 (2015).

43. Siepel, A. et al. Evolutionarily conserved elements in vertebrate, insect, worm, and yeast genomes. Genome Res. 15, 1034–1050 (2005).

44. Pollard, K. S., Hubisz, M. J., Rosenbloom, K. R. & Siepel, A. Detection of nonneutral substitution rates on mammalian phylogenies. Genome Res. 20, 110–121 (2010).

45. Rosenberg, A. B., Patwardhan, R. P., Shendure, J. & Seelig, G. Learning the sequence determinants of alternative splicing from millions of random sequences. Cell 163, 698–711 (2015).

46. Lewis, B. P., Green, R. E. & Brenner, S. E. Evidence for the widespread coupling of alternative splicing and nonsense-mediated mRNA decay in humans. Proc. Natl. Acad. Sci. U. S. A. 100, 189–192 (2003).

47. Ke, S. et al. Quantitative evaluation of all hexamers as exonic splicing elements. Genome Res. 21, 1360–1374 (2011).

48. Julien, P., Miñana, B., Baeza-Centurion, P., Valcárcel, J. & Lehner, B. The complete local genotype–phenotype landscape for the alternative splicing of a human exon. Nat. Commun. 7, 11558 (2016).

49. Adamson, S. I., Zhan, L. & Graveley, B. R. High-Throughput Identification of Genetic Variation Impact on pre-mRNA Splicing Efficiency. (2017). doi:10.1101/191122

50. Gasperini, M., Starita, L. & Shendure, J. The power of multiplexed functional analysis of genetic variants. Nat. Protoc. 11, 1782–1787 (2016).

51. MacArthur, D. G. et al. Guidelines for investigating causality of sequence variants in human disease. Nature 508, 469–476 (2014).

52. Yeo, G. & Burge, C. B. Maximum entropy modeling of short sequence motifs with applications to RNA splicing signals. J. Comput. Biol. 11, 377–394 (2004).

53. Wu, X. & Bartel, D. P. kpLogo: positional k-mer analysis reveals hidden specificity in biological sequences. Nucleic Acids Res. (2017). doi:10.1093/nar/gkx323

54. Aken, B. L. et al. The Ensembl gene annotation system. Database 2016, (2016).

55. Lek, M. et al. Analysis of protein-coding genetic variation in 60,706 humans. Nature 536, 285–291 (2016).

56. Duportet, X. et al. A platform for rapid prototyping of synthetic gene networks in mammalian cells. Nucleic Acids Res. 42, 13440–13451 (2014).

57. Website. Available at: Bushnell, B. BBMap:BBMap short read aligner, and other bioinformatic tools https://sourceforge.net/projects/bbmap/. (Accessed: 22nd September 2017)

58. McLaren, W. et al. The Ensembl Variant Effect Predictor. Genome Biol. 17, 122 (2016).

59. Harrow, J. et al. GENCODE: the reference human genome annotation for The ENCODE Project. Genome Res. 22, 1760–1774 (2012).

60. Quinlan, A. R. BEDTools: The Swiss-Army Tool for Genome Feature Analysis. Curr. Protoc. Bioinformatics 47, |p11.12.1–34 (2014).

61. Wu, X. & Bartel, D. P. kpLogo: positional k-mer analysis reveals hidden specificity in biological sequences. Nucleic Acids Res. (2017). doi:10.1093/nar/gkx323

62. Xiong, H. Y. et al. RNA splicing. The human splicing code reveals new insights into the genetic determinants of disease. Science 347, 1254806 (2015).

63. Siepel, A. et al. Evolutionarily conserved elements in vertebrate, insect, worm, and yeast genomes. Genome Res. 15, 1034–1050 (2005).

